# *An Aedes aegypti* seryl-tRNA synthetase paralog controls bacteroidetes growth in the midgut

**DOI:** 10.1101/2022.08.25.505225

**Authors:** Gilbert de O. Silveira, Octávio A. C. Talyuli, Ana Beatriz Walter-Nuno, Ana Crnković, Ana C. P. Gandara, Alessandro Gaviraghi, Vanessa Bottino-Rojas, Dieter Söll, Carla Polycarpo

**Affiliations:** Instituto de Bioquímica Médica Leopoldo de Meis, Universidade Federal do Rio de Janeiro, Rio de Janeiro, Brasil; Instituto Nacional de Ciência e Tecnologia em Entomologia Molecular (INCT-EM), Brasil; Department of Molecular Biophysics and Biochemistry, Yale University, New Haven, CT, USA; Department of Chemistry, Yale University, New Haven, CT, USA

**Keywords:** Serine-tRNA ligase paralog, Aedes, Dual oxidase, bacteroidetes, Zika virus

## Abstract

Insect gut microbiota plays important roles in host physiology, such as nutrition, digestion, development, fertility, and immunity. We have found that in the intestine of *Aedes aegypti*, SLIMP (seryl-tRNA synthetase like insect mitochondrial protein) knockdown followed by a blood meal promotes dysbiosis, characterized by the overgrowth of a specific bacterial phylum, Bacteroidetes. In turn, the latter decreased both infection rates and Zika virus prevalence in the mosquitoes. Previous work in *Drosophila melanogaster* showed that SLIMP is involved in protein synthesis and mitochondrial respiration in a network directly coupled to mtDNA levels. There are no other reports on this enzyme and its function in other insect species. Our work expands the knowledge of the role of these SerRS paralogs. We show that *A. aegypti* SLIMP (AaeSLIMP) clusters with SLIMPs of the Nematocera sub-order, which have lost both the tRNA binding domain and active site residues, rendering them unable to activate amino acids and aminoacylate tRNAs. Knockdown of AaeSLIMP did not significantly influence the mosquitoes’ survival, oviposition, or eclosion. It also neither affected midgut cell respiration nor mitochondrial ROS production. However, it caused dysbiosis, which led to the activation of Dual oxidase and resulted in increased midgut ROS levels. Our data indicate that the intestinal microbiota can be controlled in a blood-feeding vector by a novel, unprecedent mechanism, impacting also mosquito vectorial competence towards zika virus and possibly other pathogens as well.

**Author Summary:** Aminoacyl-tRNA synthetases (aaRS) are a family of ubiquitous enzymes responsible for the attachment of specific amino acids to their cognate tRNAs. During evolution some aaRS acquired new domains and/or suffered gene duplications, resulting in the improvement and expansion of their functions some of them being specific to a group of organisms. A paralog of seryl-tRNA synthetase restricted to the class Insecta (SLIMP) is found in Arthropoda. Our goal was to explore the role of SLIMP in the female mosquito Aedes aegypti using RNA interference. We showed that A. aegypti SLIMP (AaeSLIMP) gene expression is up-regulated upon blood feeding through a heme-dependent signaling. Although AaeSLIMP knockdown neither impacted the mosquito survival nor oviposition, it provoked ROS levels augmentation in the midgut via Dual Oxidase activity in order to control the increase in the intestinal native microbiota, specifically bacteria of the Bacteroidetes phylum. Although dysbiosis can result from mitochondrial impairment, this is the first time that the absence of a mitochondrial enzyme is linked to intestinal microbiota without any visible effects in mitochondrial respiration and mitochondrial ROS production. Furthermore, Zika Virus infection of AaeSLIMP silenced mosquitoes is decreased when comparing to control, meaning that Bacteroidetes overgrowth may be protecting the female mosquito. Our data indicate that the intestinal microbiota can be controlled in a blood-feeding vector by a novel, unprecedent mechanism, impacting also mosquito vectorial competence towards zika virus and possibly other pathogens as well.

## Introduction

Aminoacyl-tRNA synthetases (aaRSs) catalyze the first step of protein synthesis, tRNA aminoacylation. The aminoacyl-tRNAs formed can serve as the mRNA translators in the ribosome. There are at least one aaRS for each amino acid. Throughout evolution, gene duplication events and the loss or gain of domains/insertions have led to the expansion of aaRSs functions, some of which are directly associated with the aminoacylation catalytic site and some to the appended domains. Among the non-canonical functions of aaRSs, they can act as cytokines, take part in transcription and translational regulation, and have other enzymatic activities [reviewed in 1].

Seryl-tRNA synthetases (SerRSs) are dimeric enzymes that catalyze the aminoacylation of tRNASer with its cognate amino acid, serine. They belong to the class II aaRSs, and their catalytic core is composed of an antiparallel β-sheet flanked by α-helices harboring three motifs [2,3]. Motif 1 is related to the dimer interface and is far from the active site, whereas motifs 2 and 3 are closer together and are involved in ATP, amino acid, and tRNA acceptor stem binding [4]. In general, the N-terminally placed tRNA binding domain has a coiled-coil structure. Although many metazoan aaRSs have only one genomic copy that can act both in the cytoplasm and mitochondria, SerRSs are among a few that present compartment-specific copies. The compartment-specific SerRSs reflect the fact that the organellar tRNA^Ser^ molecules have atypical structures that must be recognized by a specific SerRS [5]. Moonlighting SerRSs were shown to exist in vertebrates. The non-canonical activity is dependent on the presence of a unique domain (UNE-S) that harbors a nuclear localization signal. This domain directs the SerRS to the nucleus, where it attenuates the vascular endothelial growth factor expression, an activity essential for vascular development [6].

In addition to the cytoplasmic and mitochondrial genes found in eukaryotes, other examples of putative SerRS proteins have been described in various organisms. In *Streptomyces sp*., SerRS homologs participate in antibiotic production [7]. These SerRS-like proteins have diverged significantly from canonical SerRSs but kept the aminoacylation capacity. In 1994, BirA, a SerRS paralog present in bacteria, was reported. It is a bifunctional protein that acts as (a) a biotin ligase and as (b) a biotin transcriptional regulator, binding to its operon and using biotin as a corepressor [8]. In some bacteria, the N-terminally truncated SerRS homologs lack the tRNA aminoacylation activity but transfer serine (and other amino acids) to the phosphopantetheine prosthetic group putative carrier proteins [9]. In 2010, a new SerRS paralog (SLIMP) was reported in some members of the Arthropoda phylum. This work showed that this paralog is crucial for the survival of *D. melanogaster*, and plays an essential role in protecting flies against oxidative stress [10]. In 2019, the same group published a study showing that SLIMP from *D. melanogaster* (DmelSLIMP) regulates mitochondrial protein synthesis and DNA replication via its interaction with mitochondrial protease LON, stimulating proteolysis of the DNA-binding protein TFAM and thus preventing mitochondrial DNA accumulation [11].

*Aedes aegypti* mosquito is a vector of Dengue, Chikungunya, Zika, and Yellow fever viruses, which cause very debilitating diseases. Blood-feeding mosquitoes are especially important because the diseases they transmit are among the leading causes of mortality and morbidity in the world [12]. These insects share the capacity to ingest massive quantities of blood in a single meal. Once blood proteins are digested in the midgut, substantial amounts of heme are released. Although heme is an essential molecule for biochemical processes such as cell signaling, respiration, and oxygen transport, an imbalance of its free amounts in cells can put tissues at risk of oxidative damage [13–15]. Mosquitoes have evolved protective mechanisms to counteract the harmful effects of free heme released after the blood meal. These mechanisms range from aggregation and degradation of heme to the induction of antioxidant defenses and detoxification genes after blood feeding, thereby reducing the amount of reactive oxygen species that can be generated [reviewed in 16].

The abundance of nutrients, together with the limitation of an oxidative burst, leads to a permissive state for the growth of the microbiota, which can increase 100-to 1000-fold [17], with a concomitant decrease in diversity [18,19]. The gut microbiota plays a vital role in the maintenance of insect metabolism. In mosquitoes, red blood cells are lysed by intestinal microbiota to accelerate blood digestion [20]. Although it has been shown that microbiota is essential for micronutrient production in several insects [21], its composition may vary among different mosquitoes. However, some bacterial groups are more abundant than others and are mainly composed of Gram-negative, oxygen-adapted bacteria belonging to Proteobacteria, Actinobacteria, Firmicutes, and Bacteroidetes [22–24].

Given the role of SLIMP in protecting against oxidative species and maintaining mitochondrial balance in flies, and considering that hematophagous insects use different mechanisms to protect themselves against pro-oxidant effects of heme, we investigated here whether *A. aegypti* SLIMP (AaeSLIMP) contributes to the maintenance of redox balance in the midgut after a blood meal. Our results show that AaeSLIMP knockdown in female *A. aegypti* mosquitoes leads to increased ROS production in the midgut. Importantly, the ROS increase is mediated by the Dual Oxidase (DuOx) and is not a result of mitochondrial ROS leakage, as in Drosophila larvae [10]. DuOx activity was triggered by the expansion of the gut bacterial load, which occurred due to the growth of a specific phylum, Bacteroidetes. Furthermore, the infection intensity and prevalence of Zika virus-infected mosquitoes were lower in those where AaeSLIMP was silenced when compared to control, demonstrating that Bacteroidetes overgrowth decreasesmosquitoes’ infection. Collectively, these findings provide evidence for an important link between SLIMP and microbiota control and add a degree of sophistication to the role of SLIMP in insect physiology, opening space for new discussion and paving the way for novel discoveries regarding the mitochondria-microbiota crosstalk.

## Results

### AaeSLIMP is localized in the mitochondria and clusters together with other Nematocera SLIMPs

The *A. aegypti* genome harbors 3 copies of SerRS genes. Two genes encode the canonical cytoplasmic and mitochondrial SerRSs; the third is a seryl-tRNA synthetase-like insect mitochondrial protein (SLIMP). Mosquitoes and flies belong to the order Diptera, which is divided into two suborders: Nematocera and Brachycera. Mosquitoes, biting midges and other flies exemplify the Nematocera, while the Brachycera include horse flies, house flies, and others. The global alignment of SLIMP sequences from Diptera shows that all SLIMPs lack the active site residues necessary for the housekeeping activity of SerRS enzymes (**Fig. S1**). However, while most SLIMPs of the Brachycera suborder, possess all SerRS sequence features, i.e., the tRNA binding domain and motifs 1, 2, and 3, the Nematocera SLIMPs and some members of the Brachycera suborder, have lost both the tRNA binding domain and motif 2. The functional consequences of the motif 2 removals can be seen in the 3D model of AaeSLIMP compared to *Bos taurus* SerRS (**Fig. 1a**).

**Fig 1.**
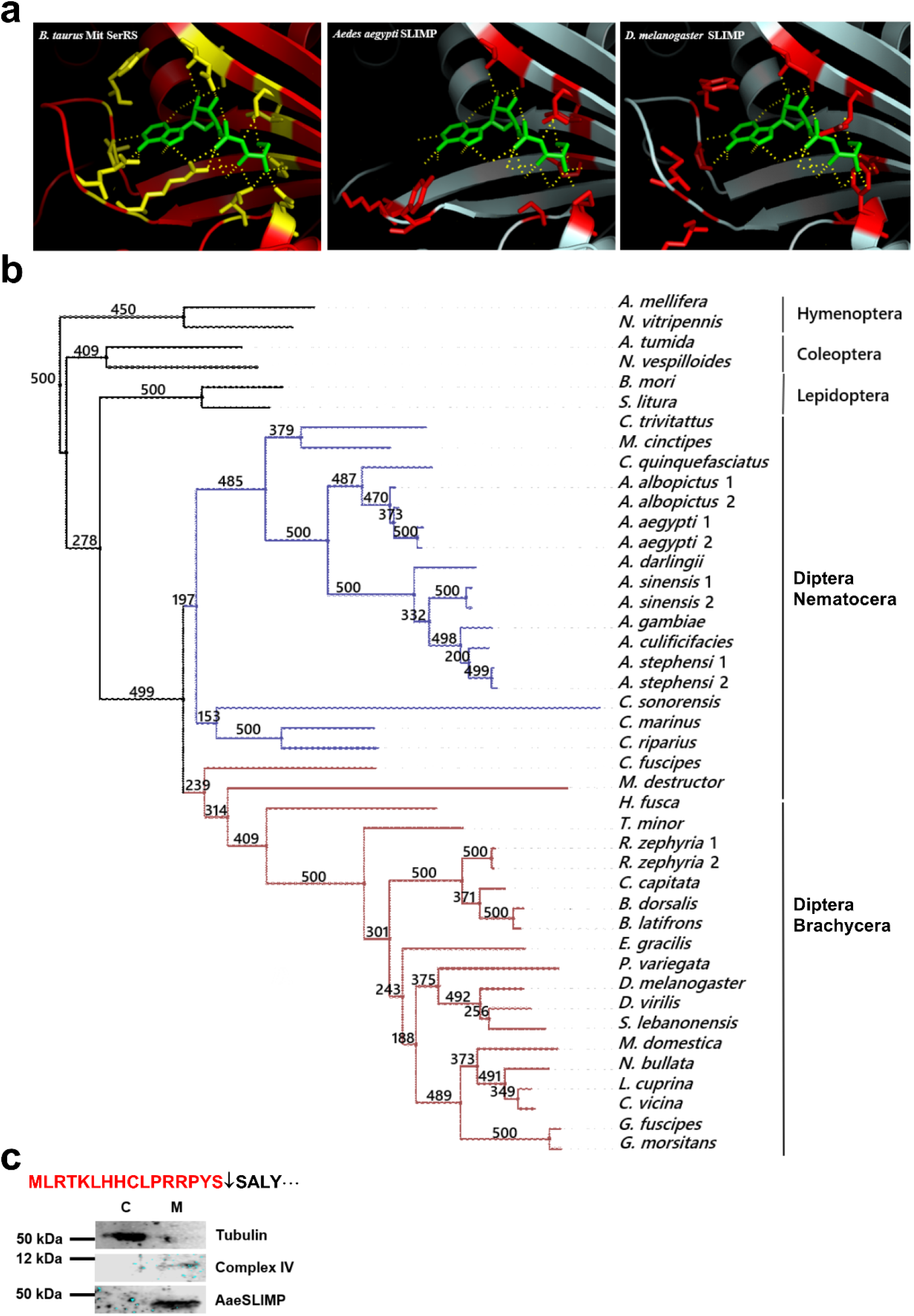
*A. aegypti* SLIMP is a mitochondrial protein that lost the SerRS motif 2. **a** Homology-based modeling of *A. aegypti* SLIMP. In green seryl-adenylate intermediate with its interaction in the active site in yellow. *Bos taurus* mitochondrial SerRS was used as the template for AaeSLIMP modeling and is represented on the left side. Essential amino acids for SerRS activity are shown in yellow (for *B. taurus* Mit SerRS) and red (for *A. aegypti* and *D. melanogaster* SLIMPs). **b** Distribuition of SLIMP in Insecta. All sequences are represented by their genus and species name. The numbers located right after each species represent the different copies of SLIMP present in their genomes. Diptera Nematocera members that did not keep motif 2 are colored in blue, and Diptera Nematocera and Brachycera that kept motif 2 are in red. Bootstrap values were obtained from 500 replicates. **c** Localization of AaeSLIMP: mitochondrial import signal predicted by MitoProt is evidenced in red from the protein N-terminus to the C-terminus and the cleavage site by an arrow pointing downwards. An enriched mitochondrial and cytoplasmic fraction from thoraxes of sugar fed female mosquitoes was used for the western blotting. Rat monoclonal anti-tubulin (1:3000 dilution) was used as cytoplasmic fraction positive control (C). Rabbit polyclonal anti-Complex IV (1:500 dilution) was used as mitochondrial fraction positive control (M). Mouse polyclonal anti-AaeSLIMP (1:2500) was used as well.

The phylogenetic tree with Diptera, Lepidoptera, Coleoptera, and Hymenoptera reflects the observed sequence differences among SLIMPs from Nematocera and Brachycera, which cluster separately with a high bootstrap value (499) and just as in the classical Arthropoda evolutionary tree (**Fig. 1b**). To confirm that AaeSLIMP is not a canonical SerRS, we cloned the AaeSLIMP gene (AAEL006938), expressed it recombinantly in a heterologous system, and purified it to perform *in vitro* studies. Like the *D. melanogaster* SLIMP, in the conditions we tested, AaeSLIMP can neither activate a mixture of amino acids nor aminoacylate total tRNA from *E. coli* or any of the *A. aegypti* tRNA^Ser^ (**Figs. S2 and S3**). Also, as for the DmelSLIMP, the MitoProtII software program [25] predicted the presence of a 15-amino acid long, N-terminal mitochondrial import peptide for AaeSLIMP (**Fig. 1c)**, localizing it to the mitochondria with a probability of 71.54%. Thus, using AaeSLIMP-specific antibodies, we performed a western blot of mitochondria-enriched and cytoplasmic fractions of *A. aegypti* thoraxes showing that AaeSLIMP is only present in the mitochondrial-enriched fraction and not in the cytosol (**Fig. 1c**).

### Heme up-regulates *A. aegypti* SLIMP gene expression

Knowing that DmelSLIMP has a role in redox balance and since hematophagous insects have evolved efficient ways to prevent ROS to increase in the gut after blood intake, we decided to examine if SLIMP had any role in *A. aegypti’s* midgut redox balance during blood digestion. AaeSLIMP’s gene and protein expression profiles were evaluated before and after a blood-feeding. We observed a statistically significant increment of AaeSLIMP gene expression 18 hours after blood ingestion (**Fig. 2a**). Additionally, AaeSLIMP protein was only detected in the protein extract from intestinal epithelium of blood-fed mosquitoes and not of sugar-fed mosquitoes, as shown in **Fig. 2b**. Knowing that heme is recognized as a signaling molecule [26,27], experiments were performed to test if it could regulate AaeSLIMP gene expression. To do so, we used an artificial diet with defined composition [28], to which we added, or omitted heme. The mosquitoes fed with the heme supplemented diet showed an increase in AaeSLIMP gene expression 18 hours after feeding (**Fig. 2c**). Notably, the same statistically significant increase of 1.3 times in AaeSLIMP expression (**Fig. 2c**) was observed when mosquitoes were fed with the natural blood meal (**Fig. 2a**), confirming our hypothesis that heme might control AaeSLIMP expression. Furthermore, an artificial diet without heme did not induce an increase in AaeSLIMP expression (**Fig. 2c**, control), proving that neither epithelium expansion nor nutritional intake triggers AaeSLIMP expression. It is well known that feeding, both with blood and an artificial diet, provides a nutritional and redox environment that favors commensal microbiota growth [17,28]. To confirm that AaeSLIMP gene upregulation expression was responsive to heme and not to bacterial growth, we treated the mosquitoes with antibiotics three days before blood-feeding to abolish gut microbiota. AaeSLIMP gene expression was up-regulated (**Fig. 2d**) even when the microbiota levels were decreased to the minimum (**Fig. 2e**), confirming that heme is responsible for the upregulation of AaeSLIMP gene expression.

**Fig 2.**
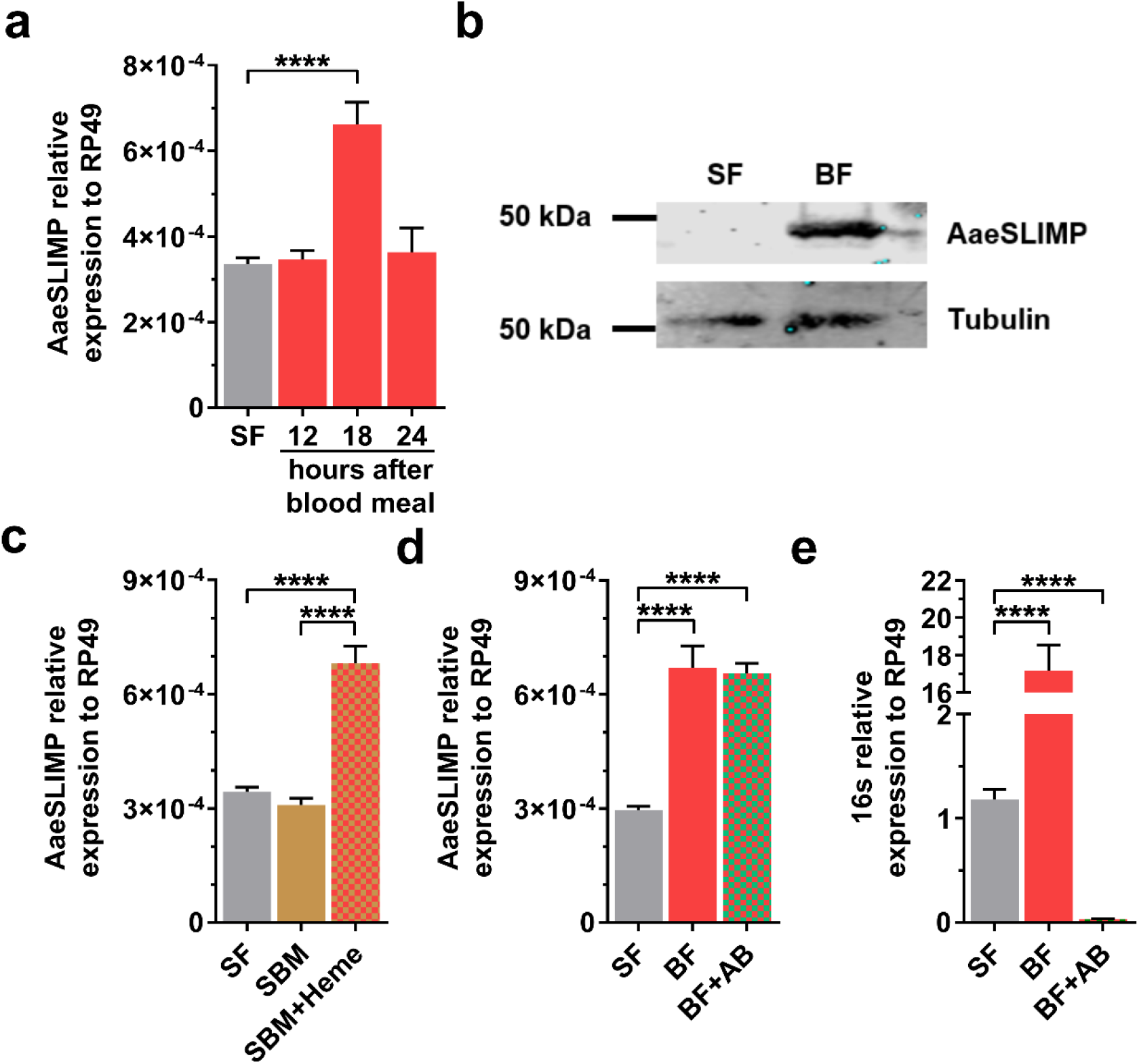
AaeSLIMP is up-regulated by heme from blood digestion in the midgut of *A. aegypti* mosquitoes. AaeSLIMP gene expression in the midgut of sugar fed (SF) and Blood Fed (BF) mosquitoes. **a** Time course of midgut AaeSLIMP gene expression before (SF) and 12, 18 and 24 hours after a blood meal. **b** Western blotting of midguts from SF and 18 hours blood-fed (BF) mosquitoes. **c** AaeSLIMP gene expression in the midgut of mosquitoes fed with Supplemented Blood Meal (SBM) with or without 50 mM heme. **d, e** AaeSLIMP gene expression and bacterial ribosomal 16S RNA expression in the midgut of mosquitoes pre-treated with streptomycin and penicillin for three days before blood-feeding to diminish microbiota levels (BF+AB). Rp49 was used as endogenous control. The average from at least 3 independent experiments is shown. Error bars are indicated. (**** p<0.0001; by Student’s t-tes t).

### AaeSLIMP knockdown promotes an increase in ROS levels via Dual Oxidase activity

To test for the AaeSLIMP function in mosquito physiology, we used RNAi to decrease the RNA levels of AaeSLIMP transcripts (by injecting a double-stranded RNA (dsRNA) that will specifically target AaeSLIMP transcripts (dsAaeSLIMP). We achieved a transcriptional reduction greater than 70% in all organs tested 18 hours after the blood-feeding (**Fig. S4 a, c, and d**). Western blotting confirmed the knockdown against the dsAaeSLIMP midgut cells protein extract, where there was no detectable AaeSLIMP protein (**Fig. S4b**). The AaeSLIMP knockdown resulted in mild to any impact on the fitness and reproductive costs of the mosquitoes (**Fig. S4 e-g**).

Considering that the knockout of DmelSLIMP resulted in increased ROS levels, we tested the hypothesis that AaeSLIMP also affected the redox environment. Thereupon, we measured intestinal intracellular ROS levels by two different approaches: (1) using dihydroethidium fluorophore (DHE), a non-specific redox-sensitive probe, and (2) Amplex-Red, a hydrogen peroxide-sensitive probe. We observed that AaeSLIMP knockdown led to an increase of about 50% in ROS levels in the midgut cells (**Figs. 3a and 3b**), in agreement with the effect of DmelSLIMP knockdown in flies. Therefore, we also explored AaeSLIMP knockdown effects in mitochondrial content and/or respiration parameters because of its probable mitochondrial localization. Against our expectations, AaeSLIMP knockdown did not lead to any changes in the midgut epithelial mitochondrial respiration rates as measured by the high-resolution respirometry (**Fig. 3c**). In fact, OXPHOS (oxygen consumption coupled to oxidative phosphorylation), maximum uncoupled respiration, and cytochrome *c* oxidase activity were not affected by AaeSLIMP’s knockdown (**Fig. 3c** and **Fig. S5**). Citrate synthase activity assay was used to determine mitochondrial content, which was not altered in dsSLIMP mosquitoes (**Fig. 3d**).

**Fig 3.**
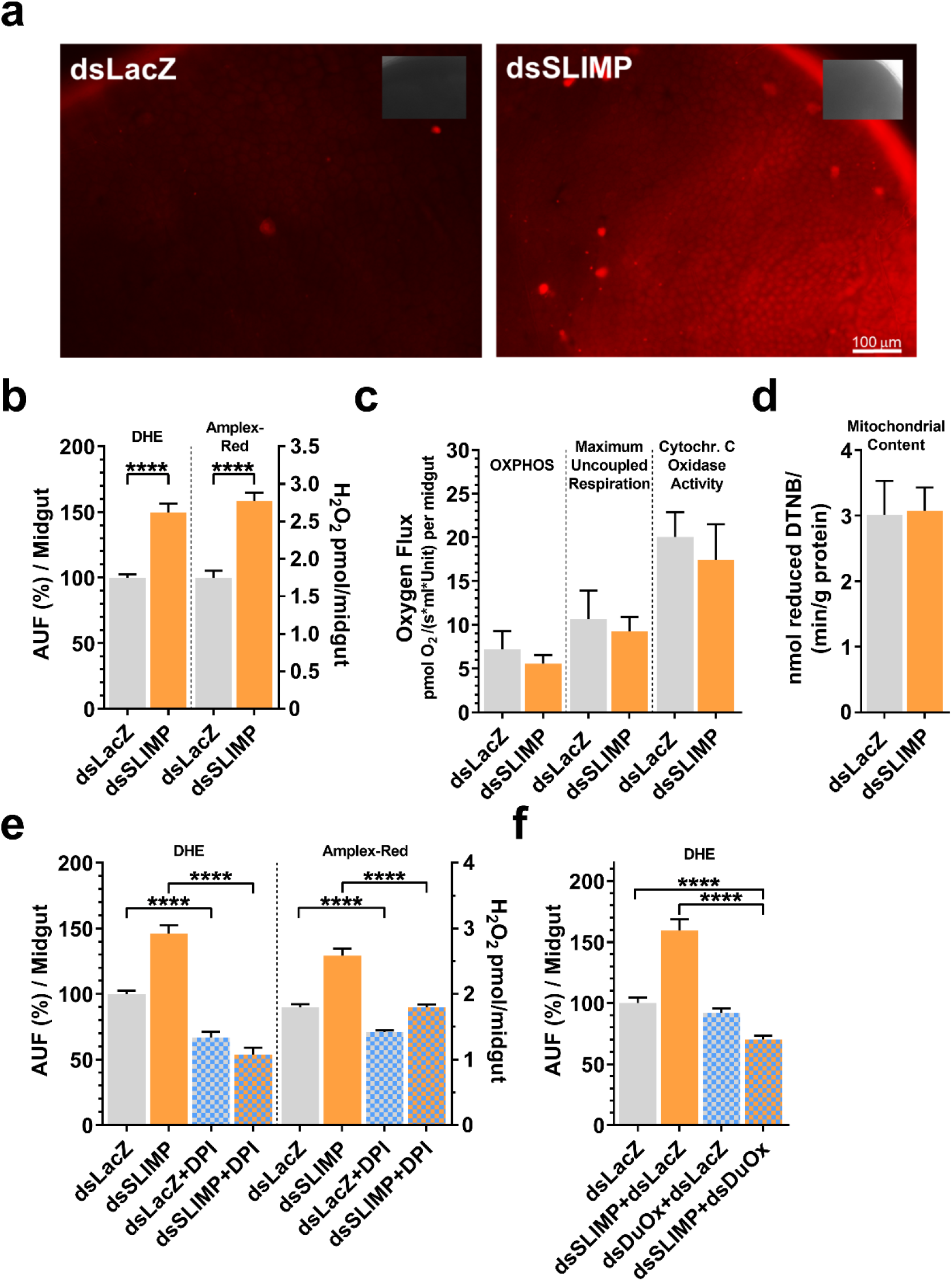
AaeSLIMP knockdown results in increased ROS production by Dual Oxidase enzyme. **a** Dissected midguts from 18 hours blood fed dsLacZ (a control dsRNA) or dsSLIMP injected mosquitoes were incubated with DHE. 100 µm scale bar. Inset: Differential interference contrast (DIC) pictures. **b, e** Midgut microscopy fluorescence quantification for DHE and resorufin fluorescence for Amplex-Red by fluorimetry. **c** Oxygen consumption assay performed with dissected midguts. OXPHOS: oxygen consumption coupled to oxidative phosphorylation. **d** Midgut citrate synthase activity was measured. **e** Before DHE and Amplex-Red incubation, dissected midguts were incubated with 10 mM diphenylene iodonium (DPI), a known inhibitor of NADPH oxidases. **f** dsRNA targeting Dual Oxidase (DuOx) was injected female mosquitoes and ROS levels were quantified by DHE. Error bars indicated. (**** p<0.0001; by Student’s t-test).

It is well known that the NADPH oxidase enzymes (DuOx, Nox4-art, and Nox5) can increase ROS production under certain conditions [17,29–32]. Since SLIMP knockdown exerts no effect on mitochondrial metabolism, we explored the role of these enzymes in the redox-altering phenotype we observed. We first checked if AaeSLIMP’s knockdown could increase DuOx, Nox4-art, and/or Nox5 enzyme and the DuOx transcriptional regulator (DuOxA) mRNAs expression. Although none of the genes tested were transcriptionally modulated after AaeSLIMP silencing (**Fig. S6a**), some are mainly regulated at the enzyme activity level for most tested animal models [33]. Hence, dissected midgut epithelium was treated with diphenylene iodonium (DPI), a known inhibitor of flavin-utilizing oxidases, and then ROS levels were measured by DHE and Amplex Red assays. Both assays showed that ROS levels in DPI-treated midguts were lower than in control (midguts of AaeSLIMP silenced mosquitoes not treated with DPI), demonstrating that flavoenzyme activity might be related to the phenotype of ROS levels rise seen after AaeSLIMP knockdown (**Fig. 3e**). Considering that DuOx is the major ROS-producing flavoenzyme present in the membranes of epithelial midgut cells in insects [17,34], we performed a double knockdown experiment to diminish mRNA levels of both AaeSLIMP and DuOx. The mRNA levels of DuOx after its silencing decreased 70-80% (**Fig. S6b**) and the midgut ROS levels in the double knocked down mosquitoes (dsSLIMP and dsDuOx) decreased 2.3 times when compared to AaeSLIMP knockdown alone (**Fig. 3f**). Thus, DuOx is the enzyme responsible for the ROS levels augmentation in the midgut cells of AaeSLIMP silenced mosquitoes.

### Midgut increased ROS levels in AaeSLIMP silenced mosquitoes are induced by microbiota dysbiosis

The DuOx-ROS system plays multiple roles in shaping the dynamic microbiome in insects, as reported for *Drosophila* and mosquitoes [17,35,36], where the microbiota overgrowth can reach 100-to 1000-fold after a blood meal [17,19]. To investigate if ROS production via DuOx affected microbiota growth in the mosquitoes’ intestines after AaeSLIMP silencing, we quantified total microbiota by RT-qPCR using 16S ribosomal subunit universal oligonucleotides. Eighteen hours after a blood meal, there was an increase of 4.2 times in 16S gene expression in the mosquitoes that had AaeSLIMP silenced (**Fig. 4a**). Using specific oligonucleotides for five different phyla (Supplementary Table 1) [37], we noticed that AaeSLIMP knockdown was causing a fivefold increase in the bacteria belonging to the Bacteroidetes phylum (**Fig. 4b**). To rule out the possibility that AaeSLIMP knockdown promotes down-regulation of an immune mediator, consequently leading to microbiota and ROS levels augmentation, we performed qPCR analysis for antimicrobial peptides as a readout of immune activation. However, we did not observe any significant changes in their expression (**Fig. S7a**).

**Fig 4.**
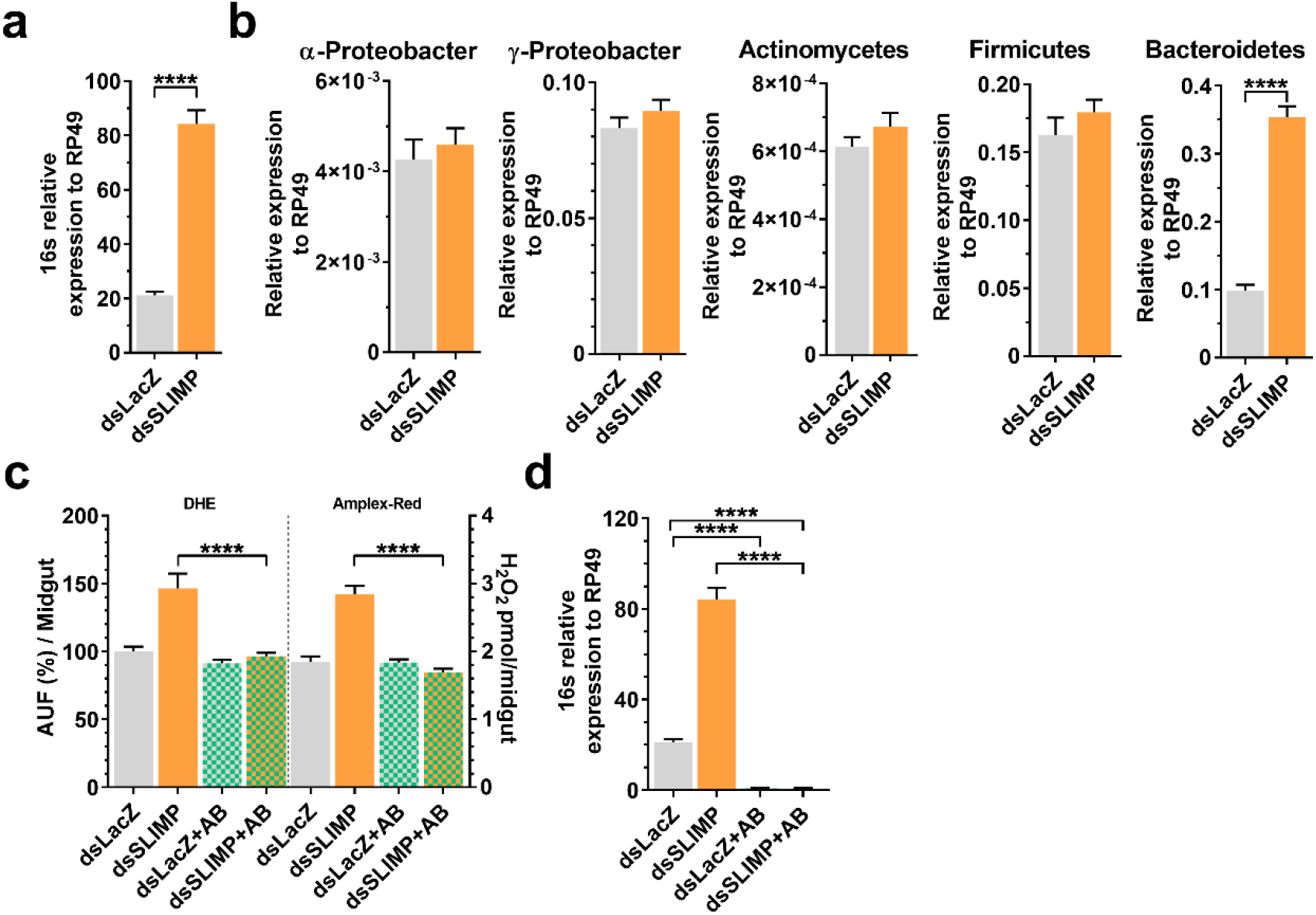
AaeSLIMP knockdown increases ROS levels because of microbiota proliferation. **a, b, d** RT-qPCR from 18 hours blood fed dissected midguts to evaluate expansion of culture-independent (16S) and phylum specific microbiota (α-and γ-Proteobacter, Actinomycetes, Firmicutes and Bacteroidetes). **c, d** Insects were pre-treated with streptomycin and penicillin for three days before dsRNA injection (+AB). **c** Dissected midguts from blood fed mosquitoes were incubated with DHE or Amplex-Red to measure ROS levels. Error bars indicated. (**** p<0.0001; by Student’s t-test).

AaeSLIMP knockdown promotes both ROS levels and microbiota increase. Since none of the immune effectors were involved in microbiota increase, we considered whether AaeSLIMP silencing would lead to midgut dysbiosis and, thus, increase DuOx’s activity. To test if the midgut dysbiosis was responsible for ROS increase, we first depleted the mosquito microbiota by feeding them with antibiotics before the blood meal and checked the ROS levels in AaeSLIMP silenced mosquitoes. **Fig. 4c** shows that the ROS levels do not increase in the midgut of dsSLIMP microbiota-depleted mosquitoes. We confirmed the microbiota levels were decreased after antibiotics treatment by analyzing the 16S transcription levels in both groups treated with antibiotics (dsLacZ+AB and dsSLIMP+AB) (**Fig. 4d**).

With those results in hand, we diminished the ROS levels in dsSLIMP mosquitoes by feeding them with ascorbate (ASC) supplemented blood meal (**Fig. 5a**). We observed that the antioxidant treatment not only decreases midgut ROS production (**Fig. 5a**), but it also allows bacteria to grow even better in dsSLIMP+ASC, than dsLacZ+ASC (**Fig. 5b**). Among these bacterial groups, Bacteroidetes increase 1.5 times (**Fig. 5c**). As we observed an increase of ROS production via DuOx in dsSLIMP mosquitoes (**Fig. 3g**), we knocked down together DuOx and AaeSLIMP and analyzed the 16S expression in the midgut. Mosquitoes that were silenced for AaeSLIMP only had an approximate increase of 4 times in bacterial growth, and dsDuOx only mosquitoes had an increase of 5 times. The dual-knockdown resulted in an additive effect, with around ten times more growth (**Fig. 5e**), which happened independently of the antimicrobial peptides’ genes upregulation (**Fig. S7**). When we look at Bacteroidetes, a similar trend is observed (**Fig. 5f**). Notably, the expression of other bacterial phyla (α-and γ-Proteobacter, Actinomycetes, and Firmicutes) increases in mosquitoes submitted to ascorbate feeding and DuOx silencing, but no difference is observed in their gene expression in mosquitoes where AaeSLIMP was also silenced (**Fig. S8**). Finally, the AaeSLIMP expression is significantly increased when ascorbate is added to the blood meal (**Fig. 5d**) and when we silence AaeDuOx (**Fig. 5g**). This result indicates that in the context of a lower oxidative state in the midgut of *A. aegypti* mosquitoes, immune mediators, as well as AaeSLIMP, must be taken into account in the control of the Bacteroidetes phylum.

**Fig 5.**
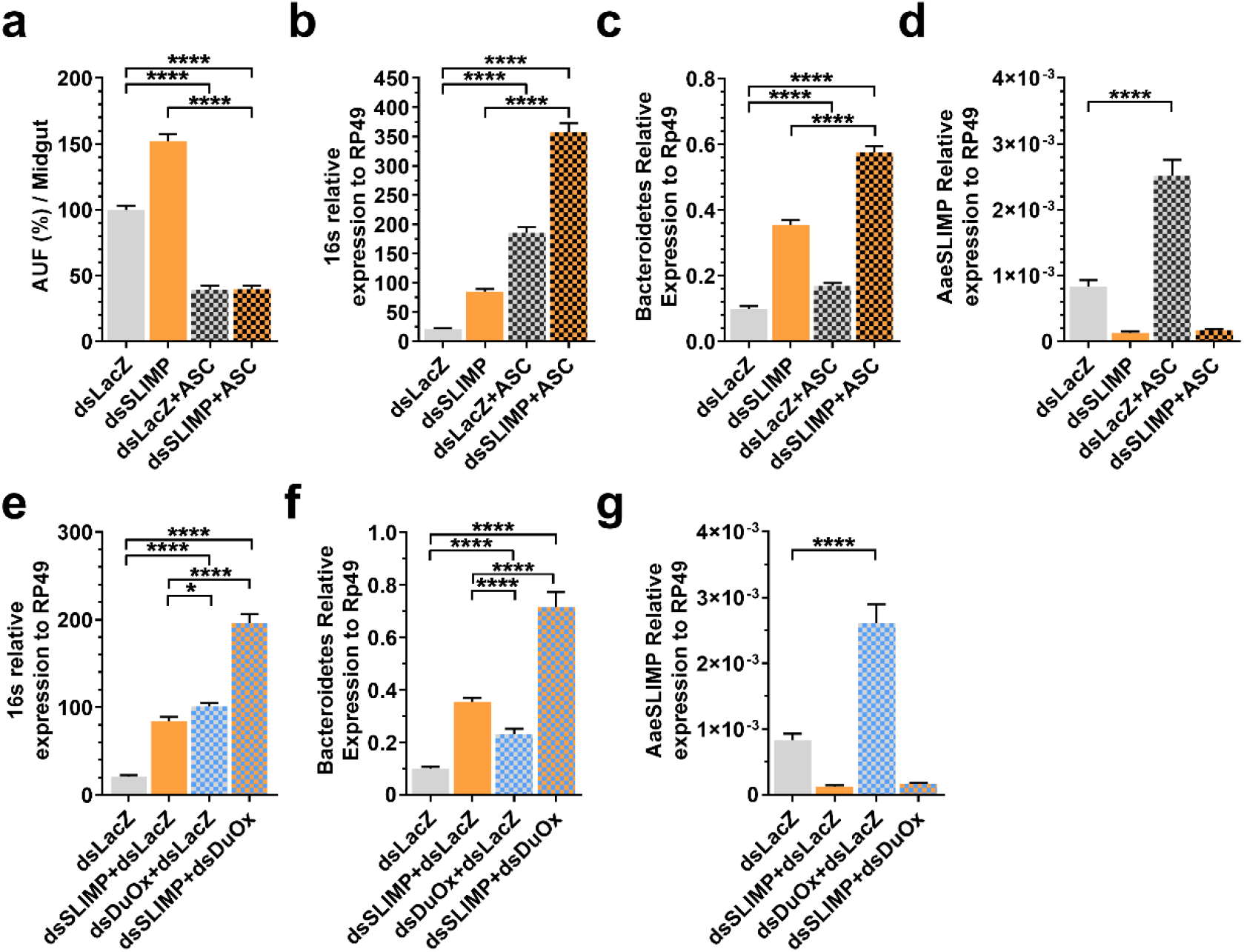
AaeSLIMP controls Bacteroidetes growth. **a** Dissected midguts from blood fed mosquitoes were incubated with DHE to measure ROS levels. **b, c, d, e, f, g** RT-qPCR from 18 hours blood fed dissected midguts to evaluate expansion of culture-independent (16S), phylum specific microbiota (Bacteroidetes) and AaeSLIMP gene expression. Ascorbate (+ASC), a known antioxidant molecule, was added to blood at a final concentration of 50 mM. Error bars indicated. (**** p<0.0001; by Student’s t-test).

### Midgut microbiota dysbiosis caused by AaeSLIMP silencing prevents the mosquito *A. aegypti* from Zika virus infection

Midgut bacteria can interact with pathogens, competing or facilitating the establishment of the infection [38]. We decided to check if the increase of the Bacteroidetes group observed after AaeSLIMP silencing would affect ZIV prevalence and infection rates in *A. aegypti*. Our results show that ZKV virus titers decreases by 3.6x in the gut of AaeSLIMP-silenced mosquitoes along with a 20% reduction in infection prevalence (**Fig. 6a and 6b**). Notably, DuOx silencing did not impact significantly the virus titers and infection prevalence in those mosquitoes when compared to control (dsLacZ). On the other hand, double-silencing of DuOx and AaeSLIMP increases the ZKV titers by 5.8x with no effect on infection prevalence when compared to control (dsLacZ), and impressively when AaeSLIMP-silenced mosquitoes are compared to double-silenced mosquitoes we see a 20x increase in ZKV titers and a 29% increase in the mosquito infection prevalence. Although these results might seem controversial, they can be explained by the fact that under the latter condition not only Bacteroidetes is augmented in the mosquitoes’ midgut and different bacteria can have different effects on microbiota, which, in this case, seems to be pro-viral. Microbiota-depleted mosquitoes (dsLacZ+AB) had ZKV titers and infection prevalence as the control mosquitoes (dsLacZ), but when comparing AaeSLIMP-silenced mosquitoes without or with antibiotics treatment, we can observe an increase of 4.8x in ZKV titers and 14% in ZKV infection prevalence in the antibiotic treated ones. Based on these results we can say that Bacteroidetes overgrowth in *Ae. aegypti* midgut has a protective effect against ZKV infection.

**Fig 6.**
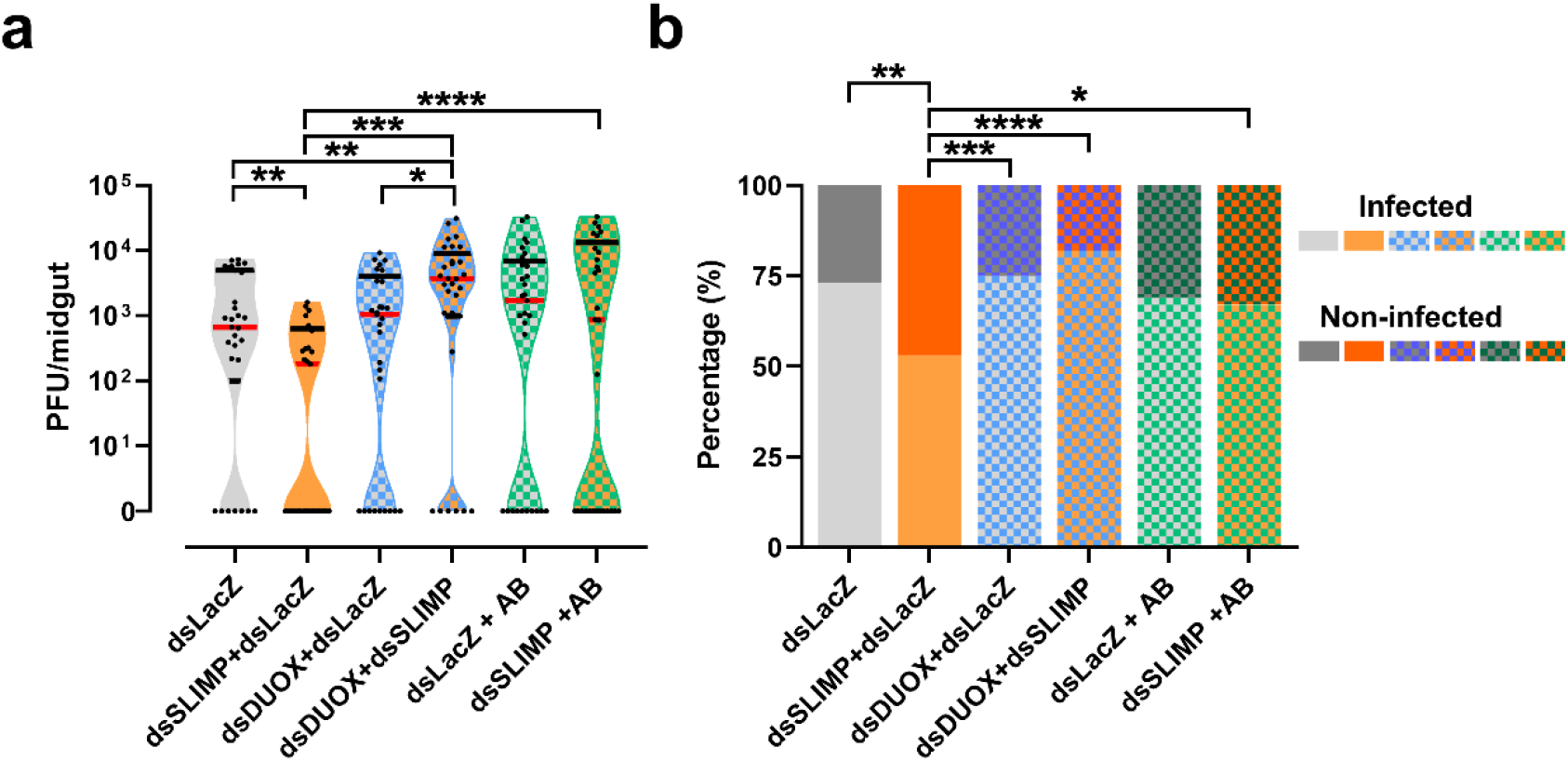
AaeSLIMP silencing impacted Zika midgut infection intensity and prevalence. **a** Females were fed blood contaminated with 10^6^ PFU/mL of Zika virus, and 7 days after feeding the number of PFU was determined in the midgut. Red and black lines represent median and quartiles, respectively. **b** The percentage of infected midguts (infection prevalence) was scored from the same set of data as in **a**. Mann-Whitney U-tests were used for infection intensity and chi-square tests were performed to determine the significance of infection prevalence analysis (*: p ≤ 0.05; **: p ≤ 0.01; ***: p ≤ 0.001; ****: p ≤ 0.0001). n=31 for all conditions tested.

## Discussion

Most paralogs of aaRS catalytic domains characterized up to date still function as ligases but often use an amino group as an amino acid acceptor instead of a hydroxyl group [reviewed in 39]. Unlike that, DmelSLIMP [10,11] and AaeSLIMP are unconventional aaRSs paralogs, as they cannot activate amino acids or hydrolyze ATP; thus, they do not have a ligase activity (shown in **Fig. S3**) [10,11]. DmelSLIMP interacts either with LON protease to control mitochondrial DNA levels or function as a heterodimer component of the canonical mitochondrial SerRS [11]. Our data on AaeSLIMP bring novel insights into these paralogs’ functions (**Fig. 7**). Diminished viability, reduced respiratory rates, and increased mitochondrial ROS production are phenotypes observed in the previous studies performed with *Drosophila* that were not observed in our experiments with midguts from *A. aegypti* blood-fed females. These results came asa surprise to us, as hematophagous insects have to survive the life-threatening conditions imposed by the pro-oxidative properties of molecules released by blood digestion and evolved a plethora of physiological adaptations to counteract this situation [16,40]. The possible antioxidant role of SLIMP in a non-hematophagous insect [10] tempted us to think it would have similar effects in *A. aegypti* mosquitoes that have fed on blood, being an additional tool against the potential oxidative imbalance in these organisms. Indeed, AaeSLIMP gene expression in midgut cells was up-regulated after a blood meal, data supported by transcriptomic analyses comparing sugar to blood-fed mosquitoes [26,27,41,42]. Also, in line with this initial hypothesis, AaeSLIMP knockdown increased the level of ROS in the midgut cells. However, we could not associate this phenomenon with mitochondrial metabolism, prompting us to evaluate other ROS-generating sources. Instead, we found the ROS increase was coming from DuOx in response to gut dysbiosis settled by the absence of AaeSLIMP. The DuOx system activity is tightly controlled at distinct levels, with MAPK p38/ATF2 controlling DuOx gene expression and intracellular calcium concentration shaping DuOx’s enzymatic activity triggered by bacterial elicitors [32,43]. None of the mosquito NOX enzymes were transcriptionally regulated in response to AaeSLIMP gene silencing (**Fig. S6a**), nor the DuOx transcriptional activator, DuOxA (**Fig. S6a**).

**Fig 7.**
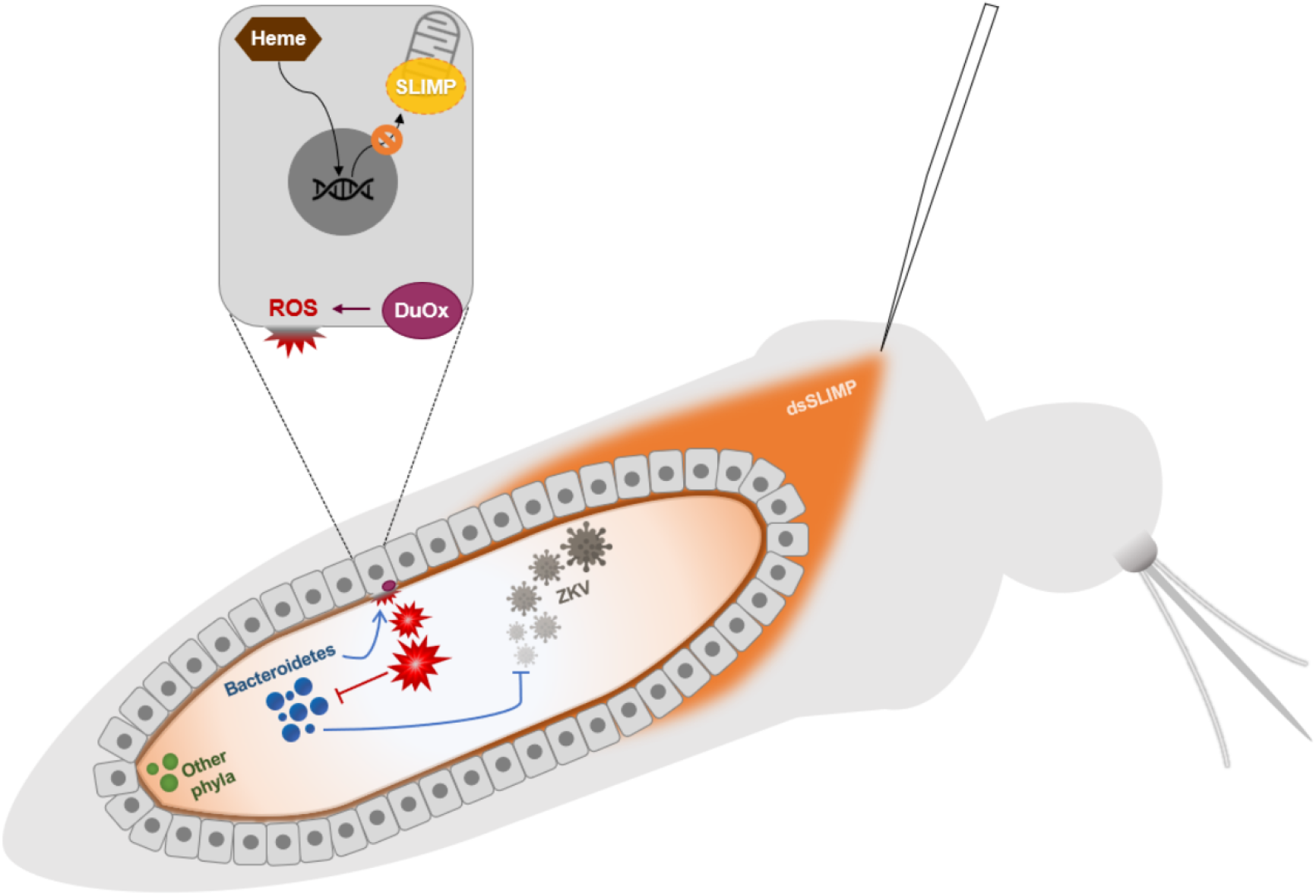
Schematic overview of AaeSLIMP effects on gut-microbiota interaction and Zika virus propagation. Blood-feeding (through heme signaling) induces expression of AaeSLIMP in the midgut. Transcriptional ablation (dsSLIMP) induces overgrowth of bacteria from the phylum Bacteroidetes and alteration of the redox state, mitochondria-independently, through activation of Dual oxidase. Bacteroidetes disbalance intervene mosquito infection by Zika virus (ZKV), all led by AaeSLIMP ablation.

Interestingly, we observed that double silencing of DuOx and SLIMP recapitulated the ROS levels to dsLacZ control, putting DuOx undoubtedly as the ROS source. *A. aegypti* blood meal decreases ROS levels in a DuOx-dependent manner and allows the proliferation of intestinal microbiota up to 1000 times [17]. Additionally, AaeSLIMP knockdown promotes an increase of 4 times in general microbiota (**Fig 4a**), even in higher ROS levels and independently of any immune effector peptides modulation (**Fig. S7a**). In turn, we showed that this DuOx-ROS production after AaeSLIMP knockdown was a response to the abnormal microbiota growth, once antibiotics-treated mosquitoes did not increase the ROS levels after SLIMP silencing (**Fig. 4c**). This observation adds another layer of discussion to the biological role of SLIMPs.

In that way, AaeSLIMP stands out as a possible missing link between ROS and commensal microbiota control in mosquitoes. In mammals, activated dynamin-related protein (Drp1) regulates gut microbiota composition by inhibiting Bacteroidetes in a ROS-dependent manner during hemorrhagic shock. However, the origin of ROS is attributed to mitochondria and not to NOX enzymes [44]. If the same microbiota effect was to be seen in *Drosophila*, one could hypothesize that SLIMP might be working in a way analogous to mammalian Drp1, by disrupting mitochondrial redox metabolism. However, our results indicate that the SLIMP effect on the microbiota is exerted by another, still elusive mechanism, that precedes the DuOx-dependent increase in intestinal ROS.

Insect intestinal microbiota plays different roles such as gut cell proliferation, nutrient digestion and supplementation and toxin catabolism [38,45–48]. Our results support the idea that this crosstalk between SLIMP and the commensal microbiota is capable to promote a very selective interference of the insect host in the microbial intestinal community, by the specific enrichment of the Bacteroidetes phylum (**Fig. 4b**). The dominant insect gut microbiome taxa belong to three phyla: Proteobacteria, Bacteroidetes, and Firmicutes. Microbes that are capable of handling oxidative stress are abundant in the midgut of different organisms. Symbiotic strains of Acetobacter possess a gene cluster related to ROS detoxification in *D. melanogaster* [49]. Another α-Proteobacteria seem to have specific responses to heme, producing a family of hemin binding proteins responsible for *Bartonella henselae* oxidative response [50,51]. As for the Bacteroidetes importance in Arthropoda, little is known. *Sulcia muelleri* is devoted to essential amino acid synthesis, whereas *Baumannia* is primarily devoted to cofactor and vitamin synthesis, both symbiont *Flavobacteriaceae* living together with *Homalodisca vitripennis* [52]. Other Bacteroidetes symbionts that are associated with grain and wood pest beetles confer desiccation resistance [53]. On a downside, some Bacteroidetes species (especially *Cardinium*) have been implicated as causative agents of reproductive incompatibility, parthenogenesis, or feminization in some arthropods [54,55]. It is interesting to note that in humans, pathological dysbiosis is primarily centered in Bacteroidetes and Firmicutes. A small number of Bacteroidetes and Firmicutes are associated with inflammatory bowel disease, especially in active inflammation regions. These bacteria produce short-chain fatty-acid metabolites, which have potent anti-inflammatory properties and may enhance epithelial barrier integrity [56].

Intestinal microbiota composition can affect directly or indirectly the ability of vectors to transmit pathogens [57,58]. Microbiota overgrowth upon insect blood feeding shapes peritrophic matrix formation which modulates viral infection through innate immune system activation [59]. However, intestinal microbiota can interfere directly in insect vectorial competence in a positively or negatively manner, depending on the bacterial species. Commensal *A. gambiae* intestinal *Enterobacter* sp. secretes ROS that kills *Plasmodium* [60], whereas *Chromobacterium* sp. inhibits viral and parasite infection in *A. aegypti* cells and *A. gambiae* mosquitoes through protease or depsipeptide synthesis [61,62]. It was already shown that *Elizabethkingia anophelis*, a bacterium of the phylum Bacteroidetes, artificially fed to *A. albopictus* reduces ZKV infection rates [63]. On the other hand, *Serratia odorifera* gut colonization increases *A. aegypti* susceptibility to Dengue virus infection [64], *Penicillium chrysogenum* fungus facilitates *A. gambiae* infection with *Plasmodium* via up-regulation of an arginine digestion enzyme preventing the production of nitric oxide, a known microbicidal radical [65]. Similarly, *Serratia marcescens* introduction in *A. aegypti* antibiotic treated mosquitoes facilitates Dengue virus infection via secretion of a protease that digests the mosquitoes’ mucus layer [66], while *Talaromyces* sp. facilitates Dengue virus infection through digestive enzymes down-regulation and trypsin reduced activity[67]. The abolishment of microbiota in mosquitoes with antibiotics has been proven to promote parasitemia of Dengue Virus and *Plasmodium* spp. in *A. aegypti* and *Anopheles gambiae* mosquitoes [68,69]. In our study, *A. aegypti* AaeSLIMP silenced mosquitoes pre-treated with antibiotics had an increase of 5x in Zika virus infection titers and 14% in infection prevalence when compared to AaeSLIMP silenced mosquitoes without antibiotics treatment (**Fig. 6a and 6b**). Although microbiota and vector competence crosstalk has been thoroughly explored a lot more needs to be understood on the mechanisms that work on these situations. Our work brings a mitochondrial enzyme to the center of this discussion for the first time.

To end, although the activity of AeSLIMP appears to contrast that of DmelSLIMP, this may not be unexpected, as *A. aegypti* and *D. melanogaster* are long evolutionary-distant insects, with different life traits. A mosquito is a blood-feeding dipteran that acquires up to 5 times its weight in blood before each reproductive cycle during its lifetime, and a fruit fly is an *ad libitum* yeast-feeding insect. These differences alone may impose a different kind of context in which the SLIMP activity evolved. In addition, the midgut responses in Drosophila have yet to be investigated. It is important to note that in *Drosophila*, ROS is produced in the gut epithelium in response to pathogenic bacteria [33,36,70]. In contrast, in mosquitoes, the release of ROS by DuOx is involved in the control of endogenous microbiota [17]. In this work, we studied the AaeSLIMP’s role in mosquito physiology and found an intriguing result, showing that there is much more to be learned about the microbiota-mitochondria crosstalk. It will be interesting to understand the preferential Bacteroidetes increase upon AaeSLIMP depletion and the role of these bacteria in the gut. Overall, our study reveals that AaeSLIMP plays a vital role in microbiota homeostasis in the mosquito gut, which affect its vectorial competence for Zika virus infections, and could perhaps affect other viral infections such as Dengue, Yellow Fever and Chikungunya, although we know that the same “immune” mechanism does not always work in the same way for the different types of pathogens [71].

## Methods

### Gene search

To search SerRS and SLIMP genes in the genomes analyzed, the PF02403 (SerRS N-terminal domain) and PF00587 (aaRS class II core domain) sequences [72] were used as queries in HMMsearch [73] using the FAT pipeline [74]. All proteins were retrieved and used as queries on BLASTp [56] against the manually curated Uniprot/SwissProt protein database [75] also using FAT.

### Phylogenetic analysis

Amino acid sequences of the proteins retrieved by our genome searches were aligned locally with MUSCLE [76], visualized, and converted to Phylip format using SeaView [77]. The maximum likelihood analysis (ML) was used to construct a phylogenetic tree with the PhyML [78] using JTT matrix [79] with default parameters. A bootstrap analysis with 500 replicates was performed to infer branch support.

### Bioinformatics Analyses

3D models were constructed by a homology-based method using three different software: Expasy-Swiss Model automatic [80], MHOLline [81], and FFPred 2.0 [82]. All models were scored according to their coverage, sequence identity, Ramachandran plot, and RMSD compared to their respective templates. The best model was analyzed with PyMOL [83], and the residues interacting with seryl-adenylate, the serylation reaction intermediate, were defined as previously reported [10] and compared to *Bos taurus* SerRS·Ser-AMP structure [5].

### Mosquitoes

*A. aegypti* (Red Eye strain) were raised in an insectary at the Federal University of Rio de Janeiro, Brazil, under a 12 h light/dark cycle at 28 °C and 70–80% relative humidity. Larvae were fed with dog chow. Adults were maintained in cages and fed using a solution of 10% sucrose *ad libitum*. Four-to seven-day-old females were used in the experiments.

Female mosquitoes were fed on blood to measure survival rate, setting the start on the survival curve. Thirty females were separated and kept in a cage (10 cm diameter x 15 cm height), and the survival was counted until the last mosquito was dead. Oviposition was performed with fully engorged females that were transferred to individual cages. The eggs were laid onto a wet piece of filter paper and counted seven days after the meal. Eclosion was measured by putting 30 eggs from individual females in water and waiting eight days to hatch.

For depletion of mosquito’s microbiota, females were fed with 10% sucrose supplemented with antibiotics -penicillin (100 U/mL) and streptomycin (100 μg/mL) (LGC biotecnologia) - for three days as previously described [17].

Mosquitoes were fed with blood directly on the rabbit’s (New Zealand strain) ear or on heparinized blood supplemented with ascorbic acid (50 mM, solubilized in 10mM phosphate buffer neutralized to pH 7 with NaOH) obtained from the rabbit’s ear vein. Mosquitoes were fed with the help of water-jacketed artificial feeders maintained at 37 °C and sealed with Parafilm membranes.

For flavoenzyme inhibition assays, the midgut dissection was carried out in a drop of PBS at room temperature. Twenty to thirty midguts were transferred to a 24-well tissue culture flask containing 1 mL of L-15 medium supplemented with 5% fetal bovine serum without antibiotics. The midguts were incubated at 25 °C for 25 minutes with 25 μM DPI (Sigma).

### Overexpression and purification of recombinant *A. aegypti* SLIMP (AAEL006938) and cytoplasmic SerRS (AAEL005037)

AaeSLIMP was cloned into pDEST17 between *Nde*I and *Bam*HI restriction sites. *A. aegypti* cytoplasmic SerRS (AaeCytSerRS) was cloned into pDEST17 between *Xho*I and *Bam*HI restriction sites. The mitochondrial import signal predicted by MitoProt [25] was not included in the AaeSLIMP coding sequence. Plasmids pDEST17 containing either AaeSLIMP or AaeCytSerRS were transformed into BL21(DE3) strain. Cells were grown at 37 °C in 2xYT medium containing 100 mg/L ampicillin. After reaching OD_600_ 0.6, cells were cooled and supplemented with 0.3 mM isopropyl β-D-1-thiogalactopyranoside (IPTG). After induction, cells were further incubated at 14 °C overnight. Cells were then harvested and resuspended in buffer A (200 mM Tris pH 8.0, 500 mM NaCl, 5 mM imidazole, 1% Triton X-100, 10% glycerol, 10 mM β-mercaptoethanol) supplemented with 2 mg/mL lysozyme. After breaking the cells by ultrasonic treatment, the soluble fraction was collected by centrifugation. After purification using Ni-NTA chromatography, eluted proteins were concentrated and stored in buffer S (10 mM Tris pH 7.5, 10 mM KCl, 10 mM MgCl_2_, 10 mM β-mercaptoethanol, 50% glycerol) at -80 °C. Immediately before the enzymatic assay, AaeSLIMP and AaeCytSerRS proteins were transferred to buffer E (50 mM Hepes pH 7.2, 50 mM KCl, 0.5 mM dithiothreitol, DTT), and their concentrations were measured according to their theoretical extinction coefficients (49300 and 58500 M^-1^ cm^-1^, respectively). **ATP-hydrolysis assay**

Enzyme’s activity was tested at 37 °C in a reaction containing 5 mM ATP, 5 mM each amino acid, 5 mM KF. 0.5 µCi of γ-P^32^ labeled ATP was used to monitor the hydrolysis. The reaction buffer used was 50 mM Hepes pH 7.2, 10 mM MgCl_2_, 50 mM KCl, 5 mM DTT. The enzyme concentration was 10 µM. *E. coli* SerRS was used in a control reaction. 2.5 µl of the reaction was taken at given times and mixed with equal volume of stop solution (0.4 M NaOAc pH 5.2, 0.1% SDS). 1.5 µl of each aliquot was spotted on a PEI-cellulose thin-layer chromatography (TLC) plate and developed in a buffer containing 1 M KH_2_PO_4_, 1 M urea. Radioactive spots of ATP, PP_i,_ and P_i_ (originating from both enzymatic and nonenzymatic ATP hydrolysis) were detected by imaging plates (Fuji Films). Imaging plates were scanned on a Molecular Dynamics Storm 860 Phosphoimager.

### Aminoacylation assay

*A. aegypti* cytoplasmic and Mitochondrial tRNA^Ser^ transcripts were searched using “tRNA^Ser^” as a keyword in Vector base (Gene set AaegL5.3) (https://www.vectorbase.org/). The genes encountered were verified by tRNAScan-SE [84]. tRNA_UGG (Cyt1) has 7 different gene copies (AAEL016067, AAEL016062, AAEL016065, AAEL016066, AAEL016287, AAEL016701 and AAEL01628), tRNA_UCU (Cyt2) has 10 different gene copies (AAEL016223, AAEL016198, AAEL016224, AAEL016225, AAEL016230, AAEL016564, AAEL016611, AAEL016612, AAEL016613 and AAEL01662), tRNA_UCA (Cyt3) has 1 single gene copy (AAEL016610), tRNA_UCA (Cyt4) has 3 gene copies (AAEL016099, AAEL016622 and AAEL016604), tRNA_AGC (Cyt5) has 5 different gene copies (AAEL016072, AAEL016073, AAEL016539, AAEL016530 and AAEL016071), tRNA_AGC (Cyt6) has 2 different gene copies (AEL016505, AAEL016642), tRNA_AGC (Cyt7) has 1 single gene copy (AAEL016318). All mitochondrial tRNAs have one gene copy: tRNA_UCA (Mit1) (AAEL018686), tRNA_AGC (Mit2) (AAEL018605), tRNA_UCU (Mit3) (AAEL016097), and tRNA_UCG (Mit4) h (AAEL016859). Sequences from all serine isoacceptors were synthesized following a T7 polymerase promoter and cloned into the pUC19 plasmid. All tRNAs that do not begin with cytosine had to be synthesized with a ribozyme sequence upstream the tRNA sequence following the T7 RNA promoter sequence to increase tRNA transcription efficiency [85].

tRNA transcription was performed with *Bst*NI digested plasmids containing each tRNA^Ser^. *In vitro* transcription was performed by the run-off method as reported before [86]. tRNA concentration was measured by spectrophotometry with NanoDrop 1000 (ThermosScientific).

Aminoacylation assays were conducted as reported before [87,88]. α-^32^P radiolabeled tRNAs were used in a 1:4 molar ratio with non-labeled tRNA. The aminoacylation reaction contained 0.1 mg/mL bovine serum albumin (BSA), 20 mM KCl, 10 mM MgCl_2_, 20 µM β-Mercaptoethanol, 5 mM ATP, 5 mM amino acid mixture containing 18 natural amino acids (tryptophan and tyrosine were omitted), 50 mM Hepes pH 7.5 and 10 µM enzyme. Aliquots were taken at different time points and kept on ice with 3 units of nuclease-P1 (Sigma). One hour after nuclease-P1 incubation, samples were applied to a TLC plate and run in a solvent containing acetic acid:1M NH_4_Cl:H_2_O 5:10:85 (v/v/v) for 2 hours. Radioactive spots of aminoacyl-adenylate (aa-AMP) and adenylate (AMP) were detected by imaging plates (Fuji Films). Imaging plates were scanned on a Molecular Dynamics Storm 860 Phosphoimager.

### AaeSLIMP expression and anti-AeSLIMP antibodies production

Because most of AaeSLIMP was not soluble, the best way to obtain enough protein for the immunization was using a protocol for extraction from the SDS-PAGE [89]. Immunization on BalB/C mouse was performed injecting two shots of 100 and 50 μg of antigen intraperitoneally, spaced by 15 days, using Freund’s Complete adjuvant in the first boost and Freund’s Incomplete Adjuvant in the second boost. Two weeks after the second shot, the blood was extracted by cardiac puncture, and the serum was kept at – 20 °C for future use.

### Western Blotting

For AaeSLIMP subcellular localization determination, cytoplasmic and mitochondrial fractions from thorax were isolated as previously reported [90]. For the sugar-fed versus blood-fed condition and AaeSLIMP knockdown, midgut from blood-fed mosquitoes (or sugar-fed whenever mentioned) was isolated, and only the epithelium was used. The tissues were lysed with RIPA buffer (20 mM Tris-HCl pH 7.5, 150 mM NaCl, 1 mM EDTA, 1 mM EGTA, 1% NP-40, 1% sodium deoxycholate). The fractions were run on a 12% SDS-PAGE, semi-dry transferred to a PVDF membrane (GEHeathcare), and a Western blot analysis was performed using anti-AaeSLIMP polyclonal antibodies generated as above (1:2500 dilution), anti-tubulin monoclonal antibodies (AbCam Ab6161) (1:8000 dilution), and Anti-VDAC polyclonal antibodies (Abcam Ab47104) (1:500 dilution). After incubation with primary antibodies for 18 hours at 4 °C, the membranes were incubated with HRP-labeled secondary antibodies followed by Millipore Immobilon ECL reagent exposition. Blots were developed using a c-Digit imaging system (LI-COR Biosciences).

### RNAi experiments

Double-stranded RNA (dsRNA) was synthesized from templates amplified from cDNA of blood-fed midgut mosquitoes using specific primers containing a T7 tail (Supplementary Table 1). The *in vitro* dsRNA transcription reaction was adapted from a tRNA transcription protocol [68]. Briefly, reactions were performed at 37 °C for 12 h in a buffer containing 40 mM Tris-HCl (pH 8.0), 22 mM MgCl_2_, 5 mM DTT, 2 mM spermidine, 0.05% BSA, 15 mM guanosine monophosphate, 7,5 mM of each nucleoside triphosphate, amplified template DNA (0.1 µg/µL) and 5 µM of T7 RNA polymerase. The transcribed dsRNA was treated with DNase at 37 °C for 30 minutes and precipitated using 1:10 (v/v) 3 M sodium acetate pH 5.2 and 1 (v/v) of isopropanol. The pellet was washed twice with 70% ethanol and then eluted in water to reach a final concentration of 3 µg/µL. Double-stranded RNA (0.4 µg) was injected into the mosquitoes’ thorax with the help of a microinjector (Drummond Scientific). A blood-meal was provided 24 hours after dsRNA injection. For dsDuOx, the dsRNA was injected 48 hours before blood intake, followed by dsSLIMP injection 24 hours before the blood meal. The LacZ gene was used as a non-related dsRNA control and was amplified from a plasmid containing a cloned LacZ fragment.

### RNA isolation and quantitative real-time PCR analysis

Total RNA was isolated from dissected midgut epithelia, thoraces, and abdomens (carcass) of females using TRIzol (Invitrogen). Complementary DNA was synthesized using the High-Capacity cDNA Reverse transcription kit (Applied Biosystems). Quantitative gene amplification (qPCR) was performed with the StepOnePlus Real-Time PCR System (Applied Biosystems) using the Power SYBRgreen PCR Master Mix (Applied Biosystems). The Comparative Ct Method [91] was used to compare RNA abundance. The *A. aegypti* ribosomal protein 49 gene (Rp49) was used as endogenous control [92]. The assessment of midgut bacterial growth was performed through qPCR of bacterial ribosomal 16S RNA. Phylum-specific qPCR was performed as mentioned [37]. All oligonucleotides’ sequences used in the qPCR assays are available in the Supplementary Material.

### *Ex vivo* ROS and mitochondria microscopy assays

The midguts dissected from the insects for microscopy were placed in L-15 culture media (Invitrogen) supplemented with 5% (v/v) fetal bovine serum and containing the fluorescent probe. The samples were incubated in the dark at 28 °C. Initially, to assess ROS levels, the midguts were incubated with a 50 μM solution of oxidant-sensitive DHE fluorophore (Invitrogen). After 20 min incubation, the midguts were washed with 0.15 M NaCl (saline solution) and immediately transferred to a glass slide for fluorescence microscopy analysis. Quantitative evaluation of fluorescence levels was performed by acquiring images under identical conditions using a 10x objective and 200 ms exposure time in each experiment. The images were acquired in a Zeiss Observer.Z1 with a Zeiss Axio Cam MrM Zeiss, and the data was analyzed using AxioVision version 4.8 software. The #15 filter set (excitation BP 546/12 nm; beam splitter FT 580 nm; emission LP 590 nm) was used for DHE labeling.

H_2_O_2_ production was assessed by monitoring resorufin fluorescence due to the oxidation of 50 μM Amplex Red (Invitrogen) in the presence of 2.0 unit/mL of commercial horseradish peroxidase (HRP) (Sigma). Eight midguts were dissected in 2% BSA in PBS and incubated at 25 °C and dim light in Amplex Red/HRP for 30 min. Fluorescence intensity was measured in the supernatant in a spectrofluorometer plate reader (SpectraMax gemini XPS; Molecular Devices) operating at excitation and emission wavelengths 530 nm and 590 nm, respectively. Background fluorescence generated as unspecific Amplex Red oxidation by the midgut in the absence of HRP was subtracted. After each experiment, a standard curve of reagent grade H_2_O_2_ (Merck) was performed.

Citrate synthase (CS) activity was assayed according to the method described by Hansen and Sidell [93]. Pools of thirty midguts were homogenized in 50 μL of saline solution. After 2 min of decantation, 40 μL of supernatant was incubated with 7.5 mM Tris buffer (pH 8.0) containing 50 μM DTNB (5,5′-dithiobis (2-nitrobenzoic acid)) (Sigma), 300 μM acetyl-CoA and 1 mM oxaloacetate.

Immediately, DTNB reduction was measured for 10 min at 412 nm. The specific activity was calculated using the reduced DTNB molar extinction coefficient (13.6 mM).

### Respirometry analyses of permeabilized midgut preparations

Respiratory activity of midgut preparations from *A. aegypti* females was performed according to methods previously established [94], with minor modifications, using a two-channel titration injection respirometer (Oxygraph-2k, Oroboros Instruments, Innsbruck, Austria). Midguts from 25 mosquitoes were dissected in an isolation buffer consisting of 250 mM sucrose, 5 mM Tris-HCl, 2 mM EGTA, 1% (w/v) fatty acid-free BSA, pH 7.4 and washed to remove all the midgut content using the same buffer. Subsequently, the midguts were placed into the O2K chamber filled with 2 mL of “respiration buffer” (120 mM KCl, 5 mM KH_2_PO_4_, 3 mM Hepes, 1 mM EGTA, 1.5 mM MgCl_2_, and 0.2% fatty acid-free BSA, pH 7.2) supplemented with 0.0025% digitonin to induce tissue permeabilization. All experiments were conducted at 27.5 °C and under continuous stirring at 750 rpm. After 5 minutes, the routine was started by adding both NAD^+^-linked substrate (10 mM pyruvate +10 mM proline) and FAD^+^-linked substrate (10 mM succinate). Afterward, the ATP synthesis was stimulated by the addition of 1 mM ADP. The oxygen consumption coupled with OXPHOS was calculated by subtracting the oxygen consumption after substrates addition from ADP-stimulated oxygen consumption rates. The maximum non-coupled respiration was induced by stepwise titration of carbonyl cyanide *p*-(trifluoromethoxy) phenylhydrazone (FCCP) to reach final concentrations of 5 µM. Finally, respiratory rates were inhibited by the addition of 2.5 µg/mL antimycin A. Cytochrome *c* oxidase activity was measured polarographically at the end of the routine of respiratory analysis using 2 mM ascorbate and 0.5 mM *N,N,N*’,*N*’-tetramethyl-*p*-phenylenediaminedihydrochloride (TMPD), as an electron-donor regenerating system. To discriminate the oxygen consumption due to cellular respiration from the self-oxidation of TMPD, 5 mM of KCN was added at the end of each experiment, and cytochrome *c* oxidase activity was considered the oxygen consumption rate cyanide sensitive.

### Virus infection and titration

Zika virus (ZKV; Gen Bank KX197192) was propagated in *Aedes albopictus* C6/36 cell line for 7 days in Leibovitz-15 medium (Gibco #41300–039) pH 7.4 supplemented with 5% fetal bovine serum, tryptose 2.9 g/L, 10 mL of 7.5% sodium bicarbonate/L; 10 mL of 2% L-glutamine/L, 1% of non-essential amino acids (Gibco #11140050) and 1% penicillin/streptomycin at 30 ºC. The cell supernatants were collected, centrifuged at 2,500g for 5 min, and stored at -70°C until use. Mosquitoes were infected with 10^6^ PFU/ml ZKV in a reconstituted blood meal made of 45% red blood cell, 45% of ZKV virus supernatant, and 10% of rabbit serum (pre-heated at 55ºC for 45 min). Four days after Zika infection, midguts were dissected and stored at -70°C in 1.5 ml polypropylene tubes containing glass beads and DMEM media supplemented with 10% of fetal bovine serum and 1% of penicillin/streptomycin. The samples were thawed and homogenized, and serially diluted in DMEM media and incubated in 24-well plates with a semi-confluent culture of Vero cells (for ZKV samples) for 1 h at 37°C and covered with DMEM 2% fetal bovine serum + 0.8% of methylcellulose (Sigma, M0512) overlay for 4 days at 37°C and 5% CO2 incubator. The plates were fixed and stained for 45 min with 1% crystal violet in ethanol/acetone 1:1 (v:v).

### Statistical analysis

All experiments were performed at least in triplicate and samples correspond to pools of 5 – 10 insects. All analyses were performed with GraphPad Prism statistical software package (Prism 8.0, GraphPad Software). Asterisks indicate significant differences (**** p<0.0001; *** p<0.001; ** =p<0.01; * =p<0.05; ns = non-significant) and the type of test used in each analysis is described in its respective figure legend.

## Statements & Declarations

### Authors’ contributions

Conceptualization: CRP; Data Curation: GOS; Formal Analysis: GOS; Funding Acquisition: CRP. and DS; Investigation: GOS, OATC, ABW-N, AC, ACPG, AG, VB-R; Methodology: GOS, ACPG, AG, VB-R; Project Administration: GOS; Resources: CRP, DS; Supervision: CRP; Visualization: GOS, VB-R, CRP; Writing – Original Draft Preparation: GOS, CRP; Writing – Review & Editing: GOS, OATC, ABW-N, AC, ACPG, AG, VB-R, DS, CRP.

### Data Availability

Not applicable.

### Ethical approval and Consent to participate

All animal care and experimental protocols were conducted according to the Committee of Evaluation of Animal Use for Research (Federal University of Rio de Janeiro, CAUAP-UFRJ) and NIH Guide for the Care and Use of Laboratory Animals (ISBN 0–309-05377-3). CAUAP-UFRJ approved the protocols under the registry #IBM115/13 to use rabbits and #IBQM118/17 to use the mouse. Dedicated technicians work in the animal facility related to rabbit and mouse husbandry under strict guidelines to ensure careful and consistent animal handling.

### Consent to participate

Not applicable.

### Consent for publication

Not applicable.

## Acknowledgments

We would like to thank all members of the Laboratory of Biochemistry of Hematophagous Arthropods at UFRJ, especially Professor Pedro Lagerblad de Oliveira and Professor Gabriela O. Paiva-Silva for critical comments on this manuscript. We would like to thank Jaciara Loredo, Monica Sales and S. R. Cássia for technical assistance.

## Authors’ Information

**Instituto de Bioquímica Médica Leopoldo de Meis, Centro de Ciências da Saúde, Universidade Federal do Rio de Janeiro, Rio de Janeiro, Brazil**.

Gilbert de Oliveira Silveira, Octávio Augusto Cunha Talyuli, Ana Beatriz Walter-Nuno, Ana Carolina Paiva Gandara, Alessandro Gaviraghi, Vanessa Bottino-Rojas, Carla Polycarpo

**Instituto Nacional de Ciência e Tecnologia em Entomologia Molecular (INCT-EM), Brazil**.

Ana Beatriz Walter-Nuno, Carla Polycarpo

**Department of Molecular Biophysics and Biochemistry, Yale University, New Haven, CT, USA**

Ana Crnković, Dieter Söll

**Department of Chemistry, Yale University, New Haven, CT, USA**

Dieter Söll

**Instituto de Química, Universidade de São Paulo, São Paulo, SP, Brazil**

Gilbert de Oliveira Silveira

**Laboratory for Molecular Biology and Nanobiotechnology, National Institute of Chemistry, Hajdrihova 19, Ljubljana, Slovenia**

Ana Crnković,

**Department of Genetics, University of Wisconsin-Madison, Madison, WI, USA**

Ana Caroline Paiva Gandara

**Morgridge Institute for Research, Madison, WI, USA**

Ana Caroline Paiva Gandara

**Department of Microbiology and Molecular Genetics, University of California, Irvine, CA, USA**

Vanessa Bottino-Rojas

